# Y-maze performance predicts refined motor learning in mice

**DOI:** 10.1101/2025.10.30.685555

**Authors:** Isak Karlsson, Thawann Malfatti, Klas Kullander, Samer Siwani, Barbara Ciralli

## Abstract

The relationships between distinct abilities and their interdependencies during memory tasks and motor learning activities are not clear. An important question is whether being proficient in memory or motor learning tasks also translates into better performance in another, similar task - reflecting potential generalization of motor learning abilities. To investigate the correlation between memory performance and motor learning, we used a combination of behavioral tasks that assess general exploratory behavior, declarative memory, and fine motor learning in female mice. For the exploratory behavior, we used the open field task, and assessed declarative memory using the novel object recognition and Y-maze. Motor performance and learning was assessed through the vertical pole and pellet reaching task. We found a positive correlation between the Y-maze and the pellet reaching task performance, where a higher exploration rate indicated a higher success ratio in the pellet reaching task. Further, leave-one-out cross-validation prediction analysis show that Y-maze performance is a robust predictor of pellet reaching performance, showing that mice pellet reaching performance can reliably be predicted from Y-maze performance. These results can be used to conduct a pre-study for challenging motor tasks that include pre-behavioral procedures on mice. Our study shows that performance in the Y-maze predicts pellet reaching performance, indicating that these tasks can be used as pre-screening for motor learning performance and activity.

## Introduction

Memory consolidation has long been recognized as a critical component of learning and adaptation [1, 2]. However, the intricate relationships between distinct cognitive abilities and their interdependencies during memory tasks and motor learning activities remain poorly understood. Motor learning is a process that refines skilled movements through practice [3], enabling adjustments to voluntary movements. This is a key function for adaptive behavior during exploration and learning, as the movements can be performed with higher speed and accuracy [4, 5]. Motor learning involves activity across multiple interconnected brain regions, including the motor cortex, basal ganglia and cerebellum [6, 7, 8].

Interestingly, neural circuits underlying previously learned tasks can be repurposed and modified to accommodate new learning, while retaining their existing memory traces [9]. This phenomenon suggests that memories from one task are not only dynamic but also context-dependent, allowing them to coexist with and influence the formation of novel memories. Specifically, experiences can serve as a foundation for subsequent learning, enabling individuals to leverage past knowledge to inform and improve their per-formance on novel tasks [9]. Furthermore, findings from motor learning studies have consistently demonstrated individual variability in the acquisition and refinement of controlled voluntary movements [10], highlighting the complex interplay between cognitive and motor processes.

Although the biological mechanisms of episodic memory and motor learning have been extensively studied, there is a gap in knowledge regarding the correlation between individual differences in these functions. Previous research has analyzed some connection between behavior tasks, although the focus was to investigate the effect an agent had on memory, not the direct correlation. A study from Kelliny et al. [11] examined the effects of streptozotocin (STZ), a diabetogenic agent, on mice behavior, and found that STZ-treated mice showed anxiety-like behavior and reduced exploratory behavior in the open field task, and simultaneously performed worse in declarative memory tasks, by investigating spontaneous alternation in the Y-maze task and recognition memory in the novel object recognition task [11]. These tasks were performed in parallel, but the study did not assess the direct correlation between them. However, the results suggest that individual performance differences across behavioral tasks may be correlated. To address this knowledge gap, we employed a combination of behavioral tasks that assess general exploratory behavior, declarative memory, and fine motor learning in mice, allowing investigation of correlations between these distinct cognitive and motor abilities.

The selection of the Y-maze, vertical pole, open field, and object recognition tasks as potential predictors of pellet reaching success was motivated by their relevance to the neural circuits involved in contextual memory and motor learning. The Y-maze performance, which depends on the integrity of hippocampal–prefrontal and corticostriatal circuits that support spatial working memory and action selection, is particularly informative, as these circuits are also critical for targeting and adjusting a pellet reach [12, 13]. Animals with poorer Y-maze alternation may struggle with spatial memory, leading to difficulties in identifying target locations or executing effective reach strategies, which could result in lower success rates in the pellet reaching task. Also, exploratory behaviors in the Y-maze can influence motor task performance by modulating attentional resources and movement efficiency during skill acquisition [14]. Similarly, the vertical pole task, which assesses postural stability, proprioceptive ability, and cerebellar–corticospinal coordination [15], directly influences skilled reaching by evaluating the motor control and coordination necessary for precise movement. Poor performance in the vertical pole task may indicate deficits in motor execution, even when spatial targeting is accurate, suggesting that these tasks may reflect shared neural substrates that underlie both memory and motor function.

A challenge in performing animal studies where the subjects must execute a task that requires learning is the fact that the process of learning is highly individualistic. When the mechanisms of learning are not the study’s focus, it is important to reduce the experimental variance caused by the individual’s ability to learn a task. In addition, by identifying and selecting animals performing well in the task, the likelihood of successful future experiments involving more invasive procedures, such as optic ferrule implantations (optogenetics) and lens implants (miniscope) increases. To achieve this, one could perform a selective study to identify individuals with a higher likelihood to learn a challenging task based on simpler tasks involving memory and motor execution, where mice performing well in the tasks are selected for further investigation. This study aimed to find pre-screening tasks for improved performance in a challenging motor task – the pellet reaching task. We selected open field, novel object recognition, Y-maze and vertical pole, due to their simpler and shorter execution and interpretation. The hypothesis was that mice that perform well in general behavior tasks that include motor coordination may also perform well in the pellet reaching task.

## Materials and methods

### Animals

All experiments were performed with the ethical permit (number 5.8.18-07526/2023) approved by the Swedish Board of Agriculture. The animals used in this study are heterozygous mice from the Chrna2-cre line crossed with C57Bl/6J wild-type mice. Chrna2-Cre mice, which were produced and bred in our own facility [16], were initially on 129SvJ/C57BL/6J mixed background and then crossed *>* 10 generations with C57BL/6J (Taconic, Denmark) mice. Offspring were genotyped in-house for the presence of the Chrna2-Cre allele. We used Chrna2-cre mice to establish a genetic background that enables precise manipulation of Chrna2+ neurons [16], a critical tool for investigating neural mechanisms underlying contextual memory [17] and motor learning [18]. While this genetic construct was not relevant to our current study, some of these mice were used in subsequent experiments using these features, all conducted after our behavioral assays were completed. At the end of the subsequent experiments, mice were deeply anesthetized with Medetomidin 1mg/kg (Dormitor Vet, Orion Pharma Animal Health) and Ketamin (Ketalar, Pfizer) 75 mg/kg i.p. and transcardially perfused with phosphate-buffered saline (PBS, Fisher Scientific, CAT 10051163), followed by 4% formaldehyde (VWR Chemicals BDH®, CAT 9713.1000).

Twelve mice, aged four months at the beginning of the experiments, were housed in a total of 4 cages, in groups of 2 to 4 individuals per cage in a temperature-controlled room with a 12-hour cycle of daylight and darkness, with access to water ad libitum. All experiments were conducted during the light phase of the 12-h light/dark cycle. Cages were lined with wood chip bedding and enriched with paper houses and nesting materials. Food was available ad libitum, unless on occasions when the animals were food-restricted. In those cases, the restriction lasted a maximum of 20 hours. All mice used in this study were female and kept at a body weight of a minimum 90% of their starting body weight. We selected only females in order to avoid observed aggression between males during habituation sessions, when 2–3 mice were placed in the arena concurrently, which could induce stress on the day of the experiment. Furthermore, literature confirms no sex-specific differences in the behavioral metrics used here for C57Cl6/J mice on the light phase of the light/dark cycle [19, 20, 21, 22].

In the open field test, all 12 mice were included. In the novel object recognition task, nine mice were included, and three mice were temporarily excluded due to an unstable object movement during recording. In the Y-maze and the vertical pole task, all 12 mice were included. Due to an observed illness, one of the mice was excluded from the pellet reaching task, and 11 were included.

On the day of testing, all animals were transported to the behavioral test room approximately 30 minutes before the start of the task, and placed onto a table separate from the experimental setup. Following this acclimatization period, each animal was individually placed into the experimental setup and subjected to the tasks. After testing, the animals were checked for their tag number and transferred to a separate cage to minimize the risk of stress transfer to other animals. Once all animals from a cage had been tested, they were returned from the separate to their original cage. This procedure was repeated until all mice from all cages were tested. The experimenter was blind to the performance of mice on previous tasks, and due to the computational nature of all analysis, the analyzer was blind to the identity of animals.

### Habituation

The habituation procedure was conducted over the course of five consecutive days. The animals were brought to the behavior room to acclimate for 30 minutes. The room’s light intensity was gradually increased to goal-light day by day, spanning from five lux initially to 80 lux at the end of the habituation process, which was the same light intensity used in the following tasks. The duration of each habituation session was five minutes, performed twice a day with a minimum of two hours of rest between the sessions. Cages used for habituation were cleaned with 10% ethanol and a paper tissue. Before each task, the animal home cages were brought to the behavioral room to acclimatize (15-30 minutes before initiating the task). For each task, mice were placed individually in the arena. To minimize stress levels in the task, we checked the animal’s identity after the trials.

### Open field

The Open-Field test is used in this study to assess locomotion and exploratory behaviors when animals are exposed to an empty arena. The task requires little animal training, and only one test session is needed. This task was mainly used to habituate the animals to the arena before novel object recognition, and to measure a baseline for general exploratory behaviors. To assess locomotion and to habituate mice to the arena, individual mice were placed in an empty round arena, measuring 46 cm in diameter. The center region of the arena was defined as a circle of 18.5 cm around the center of the arena. For easy detection by the recording device, the chosen arena was white. The recording time was five minutes, and one session was recorded for each mouse. After five minutes, the animal was brought to a new cage, and the arena was thoroughly cleaned with 10% ethanol and a paper tissue. The behavior was recorded with a JVC camera. The extracted measurements were distance moved, velocity and time spent in the center.

### Novel object recognition

This task was performed over the span of two days [using the same parameters, settings and setup as previously described; 17]). The experiment set-up consisted of a round 46 cm diameter white arena (same as open field), two identical “familiar” objects, one “novel” object, a JVC camera for recording the material, and a computer. Two similar dish brushes that had been modified and attached to a lid, to allow them to stand upright, were used as familiar objects. The novel object was a dish brush with a different shape. For testing, animals were brought to the arena one by one and spent 10 minutes exploring the objects. After 10 minutes, the animals were brought to a new cage, and the arena was thoroughly cleaned before bringing the next animal. The objects were cleaned with 10% ethanol and a paper tissue while still being placed in the arena, to ensure that the objects are in the same place for each animal.

On the first test day, the animals were exposed to two objects. The next test day, one of the objects was replaced with a novel object. The novel object was similar to the familiar objects in terms of size but easily distinguishable. The location of the objects was counter-balanced across animals. The novel object was placed on the left side for half the group, and on the right side for the other half to minimize potential side bias. Both days were recorded using a JVC camera mounted directly above the arena. For analysis, the time spent investigating the familiar vs novel object on test day was calculated. We define investigating behavior as the behavioral state where the mouse’s head is within a circular zone around the object with distance between the object edge and the zone edge approximately equal to the size of the mouse’s head. This zone captures instances where somatosensory exploration (whisker/paw contact) occurs. The ratio was calculated as the time spent investigating the novel object divided by the sum of the times investigating the familiar and novel object, then converted to percentage by multiplying by 100.

### Y-maze

The Y-maze assesses short-term spatial working (declarative) memory in mice, driven by their innate curiosity and motivation to explore new areas [23]. The task is carried out by allowing the animal to freely explore a Y-shaped arena (L: 53.5 cm; W: 5 cm; H: 10 cm) while tracking the spontaneous alternation of entries into each arm. A continuous pattern of entry into each arm characterizes an alternation. The number of alternations is the result that indicates working memory performance. For example, if the animal has explored the maze by entering arm A and then arm B, the next arm would be arm C. If the animal has a low number of correct alternations, their working memory might be impaired. A correct alternation occurs when the animal explores all three arms of the Y-maze without re-visiting any, demonstrating its working memory capabilities and preference for novel stimuli. The total number of possible alternations is given by *TA* = *n*_visits_ *− n*_arms_ *−* 1, where *n*_visits_ is the number of visits and *n*_arms_ is the number of arms (= 3). The correct alternation ratio is then calculated as 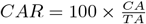, where *CA* is the number of correct alternations and *TA* is the total number of possible alternations.

For testing, the arena was elevated on the floor to minimize the risk of escapes, built directly beneath the JVC camera. The animals were brought to the behavior room and left for 30 minutes to acclimate, and then brought to the arena one by one and placed in arm B to explore the maze alone for 10 minutes. The arena was cleaned with 10% ethanol and a paper tissue. Analysis was made by calculating the number of correct alterations divided by the total number of possible alternations, multiplied by 100.

### Vertical pole

The principle of the vertical pole task is to assess motor coordination skills in mice [24]. A 50 cm high metal pole was used, covered in masking tape to make ridges that the mice can grab onto. The mice were placed at the top with their face turned upwards. The measurements for the task are the time to turn and time to descend. The time to turn describes the time it takes for the mouse to turn from being placed on top of the pole, face up, until it has turned its body 180° in alignment with the pole. The time to descend measures the time for the mouse from being placed on the pole, faced up, until it has reached the bottom of the pole [24].

Before training and testing, the animals were allowed to acclimate in the room for 30 minutes. The pole was placed in an empty cage on a table in the experimental room. The animals were trained for two days before the task, and during these sessions, they were habituated to the test setting. Each animal was trained to climb the pole by being placed halfway up, faced down, and allowed to climb down the pole with support from the experimenter as needed to reduce the risk of falling. When the animal had managed to climb down half the pole, it was instead placed at the top of the pole, faced down, and allowed to climb down freely with support if needed. After successful climbs, the mouse was placed an additional time at the top of the pole, this time faced up, to simulate the real test setting and allowed to climb down freely. This training trial was repeated until the animal was able to climb down securely. In between the training trials, the animal was left to roam the cage for 30 s.

Before the test session, a video camera (Canon Legria HF M52 10x optical zoom) was mounted on a stand and directed towards the pole, with both the top and bottom in frame. All apparatuses were cleaned with 10% ethanol and a paper tissue. The task was carried out with three consecutive test trials, with 30 s of free roam time in the experiment cage between trials. Mice performance was quantified as the mean value from the 3 consecutive trials.

### Pellet reaching

The pellet reaching task is a widely used task to assess fine motor learning and skilled forelimb use in rodents [25]. The mice were trained and habituated to the arena for four consecutive days, as previously done [26], which was sufficient since none of the animals exhibited signs of anxiety or fear, such as urinating and defecating in the chamber or attempting to escape by jumping towards the chamber ceiling. Both training sessions and test sessions lasted for 10 minutes. The first two days, mice were placed in the arena in groups of two to three individuals together. Sugar pellets (Non paraille; manufacturer: Dr. Oötger) were uniformly distributed across the arena floor prior to behavioral testing. Mice were allowed to explore the arena during initial habituation trials, with the experimenter in close proximity to acclimatize mice to the experimenter’s presence. In the third and fourth session, the animals were trained one by one, and the pellets were offered on a metal spoon through the slit instead of distributed across the arena floor.

During the test sessions, the animals were tested one by one and were allowed to take a pellet only from the spoon in a 10-minute session. For each test session, the animals were food restricted to enhance their motivation to reach for the sugar pellets. Before removing the food, each animal was weighed to verify whether they had lost more than 10% of their initial body weight. All animals used in this study maintained at least 90% of their body weight throughout the experiments. Food was removed 15 hours before the experiment was initiated and put back after no more than 20 hours. Three test days were executed, with one rest day in between test days due to the food resctriction protocol.

The task was recorded with a video camera (Canon Legria HF M52 10x optical zoom) mounted on a tripod, with the arena slit in frame. The trials were categorized into successful or failed reaches. A successful reach occurred when the animal could effectively grab the pellet and eat it. A failed reach occurred when the animal could not complete the task, either by missing, hitting, and dropping the pellet, grasping and dropping before eating, or reaching for the pellet when no pellet was available. The success ratio was calculated as the total number of successes divided by the total reaches, multiplied by 100.

### Statistical analysis

The effect of object familiarity on novel object recognition and the effect of behavioral arena on total distance moved were analyzed using one-way repeated measures ANOVA, followed by paired t-tests for post-hoc pairwise comparisons, corrected for multiple comparisons using Holm’s method. Due to the low number of data points, correlation statistics were performed using the Spearman coefficient with p-values calculated via permutation with 9999 iterations, to assess the relationships between each behavioral measure and the motor learning task. r values close to +1 measure positive correlation, meaning when one variable increases, the other variable increases linearly. Values close to 0 mean there is no correlation, and values close to −1 indicate a negative correlation, meaning when one variable increases, the other variable decreases proportionally. Since we do not investigate an integrated null hypothesis across the full correlation matrix, no multiplicity correction is required for the exploratory pairwise correlation matrix [27].

To assess the relationship between Y-maze correct alternation ratio (*X*_1_) and pellet reaching success ratio (*Y*) while controlling for distance moved and maximum alternations (**C**), we computed partial Spearman rank correlations. First, linear effects of **C** were removed from both *X*_1_ and *Y* via ordinary least squares (OLS) regression. Specifically, we fitted models:

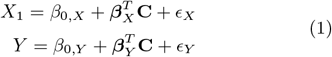

where **C** is the matrix of confounders; *β*_0,*X*_ and *β*_0,*Y*_ are the intercept terms for the models predicting *X*_1_ and *Y*, respectively; and 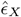 and 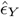 are the retained residuals. The Spearman correlation coefficient *ρ* was then calculated between these residuals. To determine statistical significance, we employed a permutation test with *N*_*perm*_ = 9999 iterations. In each iteration, the pairings between 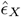 and 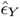 were randomly shuffled, and the Spearman statistic was recomputed to generate a null distribution. The two-tailed *p*-value was derived as the proportion of permuted statistics with absolute magnitude greater than or equal to the observed |*ρ*|.

We fitted a multiple linear regression model of the form:

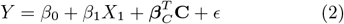

where *β*_0_ is the global intercept, *β*_1_ is the slope coefficient for the primary predictor *X*_1_, ***β***_*C*_ is the vector of coefficients for the confounders **C**, and *ϵ* is the error term; using OLS estimation. Model fit was evaluated using the coefficient of determination (*R*^2^) and adjusted *R*^2^. Significance was assessed using non-parametric bootstrapping. We performed *N*_*boot*_ = 1000 resamples with replacement, refitting the model for each iteration to extract the slope estimate for *X*_1_. The 95% confidence interval was defined by the 2.5th and 97.5th percentiles of the resulting bootstrap distribution. The slope was considered statistically significant if the confidence interval did not contain zero.

To evaluate out-of-sample predictive performance, we conducted leave-one-out cross-validation (LOOCV). In this procedure, the model was iteratively refitted on *n −* 1 mice performance, and the left-out mouse performance was predicted. Performance metrics included the mean squared error, root mean squared error, mean absolute error, and cross-validated 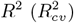, calculated as:

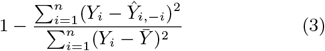

where *n* is the total number of mice, *i* is the individual mouse, 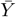 is the sample mean of *Y* across all mice, and 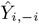 denotes the prediction for mouse *i* excluding it from the training set. A model was considered to have positive predictive validity if 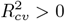.

For all violin plots, the filled area represents the data range, with a horizontal line at the median, a triangle marker at the mean and whiskers representing the standard error of the mean. For all scatter plots, circles represent individual data points for each mouse.

### Software availability

Video recordings for the open field, novel object recognition, and Y-maze tasks were obtained using EthoVision software version 15, and behavioral data tracking and extraction were performed using EthoVision software version 16. Data analysis was conducted using Scipy [28], Numpy [29] and SciScripts [30], while all plots were generated with Matplotlib [31]. Schematics were created using Inkscape [32]. The data analysis script is available online [33].

## Results

Mice were tested in the open field, novel object recognition, Y-maze, vertical pole, and pellet reaching tasks to examine whether refined motor learning skills correlate with more generalized motor performance (Fig 1A). To assess this, multiple measures were extracted from each task, including the locomotion in the open field, as a baseline for how much the individuals tend to move in the arenas (Fig 1B-L). As expected from the intrinsic drive to explore novel objects compared to familiar ones [34], most mice spent more time investigating the new object than the familiar object (8 of 9 subjects: mean time = 88.94 s for the new object vs. 47.29 s for the familiar object; Fig 1E). This shows that although 25% of the session time was spent exploring the objects, animals could remember which object was replaced, with innate curiosity motivating them to investigate the new object more than the familiar one during testing. The ratio indicates that individuals spent on average double the time examining the new object compared to the familiar one (Fig 1F). These results indicate that the group is able to distinguish the novel object and perform the task adequately.

**Figure 1:**
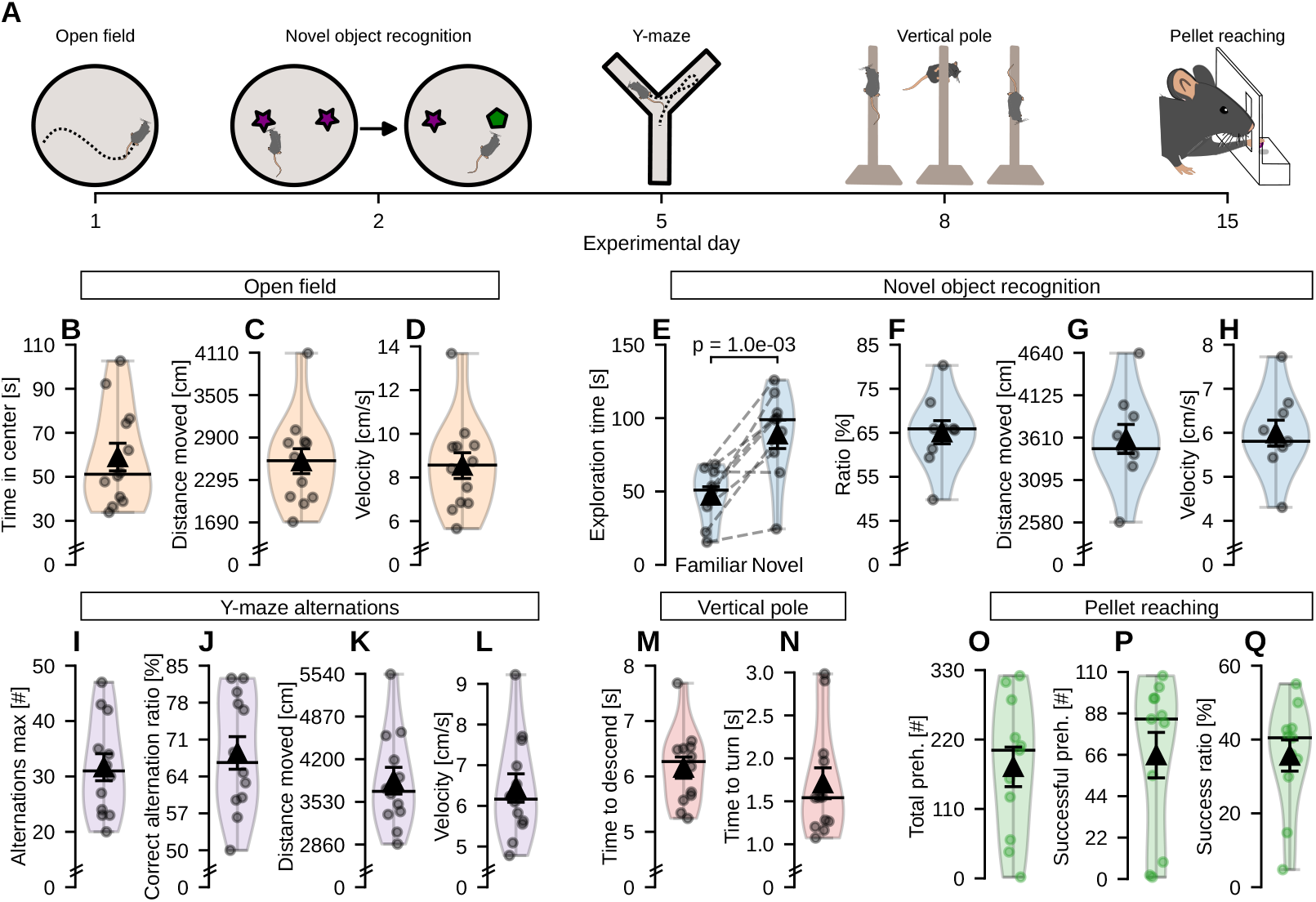
Mice performance in each behavioral task. A) Timeline of experiments, with experimental days where the tasks were initiated. B-D) Time spent in the center of the arena (B) distance moved (C), and velocity (D) during the open field test. E-F) Exploration time (E) and ratio (F) for the familiar in relation to the novel objects. G-H) Distance moved (G) and velocity (H) during the novel object recognition task. I-L) The total number of possible alternations (I) and correct alternations ratio (J), distance moved (K) and velocity (L) during the Y-maze task. M-N) Time to descend from the top of the pole to the floor (M), and time to turn on the top of the pole faced up until the animal is completely aligned with the pole faced down (N) in the vertical pole task. O-Q) Total number of reaches (O), total number of successful reaches (P) and success ratio measured in percentage (Q) for the pellet reaching task.

To assess working memory, we used the Y-maze task. Correct alternation suggests good working memory, and the mean value for the correct alternation ratio for the group was 68.5% (Fig 1I-J). The vertical pole task was used to evaluate motor coordination by measuring the mean time for animals to turn and descend from the pole (Fig 1M-N). The group mean time for the time to turn was 1.7 s, and for the time to descend, the mean time was 6.2 s. In the pellet reaching task, we evaluated how well the animals can learn a complex motor task. The mean attempts of reaches in the group was 176 times (Fig 1O), and of those, the mean value of successful reaches was 65 times (Fig 1P), giving the group a success ratio of 35% (Fig 1Q). This ratio can be interpreted as the individuals in general are active and reach for the pellets, but only a third of the time succeed in grabbing them.

The distance moved and velocity were evaluated in the open field, novel object recognition, and Y-maze arenas to verify whether any animals spent most of the time standing still in one task. Comparison of the first 5 minutes of measurements in each task (novel object recognition and Y-maze) with open field test measurements - to ensure consistent 5-minute recording duration for all tasks - indicated one outlier with greater distance traveled and faster velocity (4102.7 cm, 13.68 cm/s, 2.17 std. above the mean, 3.92 times the interquartile range above the third quartile) compared to the group mean (2709.46 cm, 9.03 cm/s), but no outliers showing reduced exploration (Fig 2A-B). Since the mouse identified as a statistical outlier did not display immobility during the tasks, no animal was excluded from further analysis. Analysis of the distance moved and velocity confirms that Y-maze tasks show significantly greater distance traveled (t-test eff. size = −1.331; p = 1.3e-02) and velocity (t-test eff. size = −1.331; p = 1.3e-02) compared to novel object recognition exploration, showing that Y-maze performance reflects active exploration rather than immobility.

**Figure 2:**
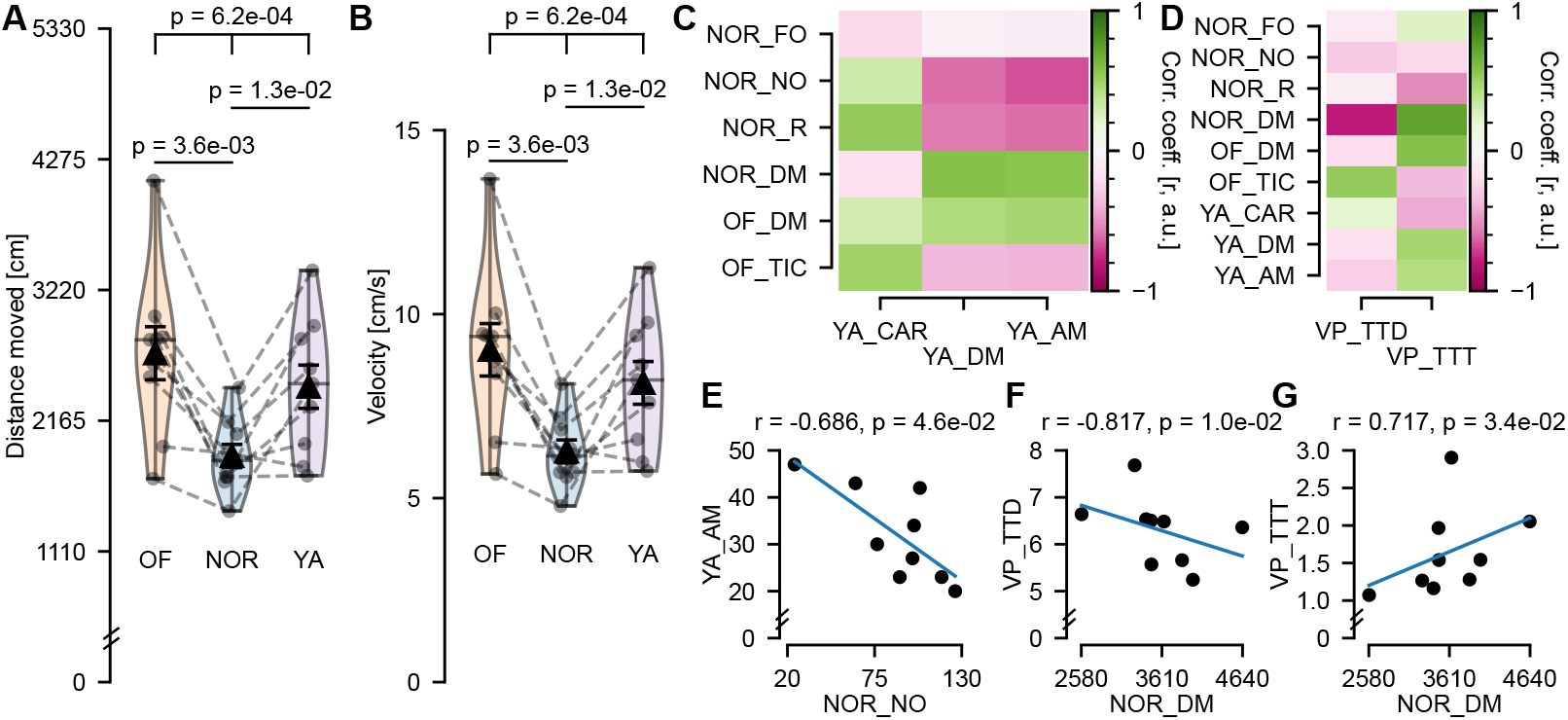
Mice with faster descending times in the vertical pole task travelled larger distances in the novel object recognition task. A-B) Distance moved (A) and velocity (B) in the open field, novel object recognition and Y-maze. C) Correlation matrix between novel object recognition and open field with y-maze measures. D) Correlation matrix of measures from novel object recognition, open field, and Y-maze with vertical pole measures. E) Correlation observed between the total number of possible alternations in the Y-maze and time exploring the novel object in the novel object recognition. F-G) Correlations observed between traveled distance in the novel object recognition task and time to descend (F) and time to turn (G) in the vertical pole task. Abbreviations: NOR (Novel Object Recognition); OF (Open Field); VP (Vertical Pole); YA (Y-maze); DM (Distance Moved); FO (Familiar Object); NO (Novel Object); R (Ratio); TIC (Time In Center); TTD (Time To Descend); TTT (Time To Turn); AM (Alternations Maximum - total number of possible alternations); CAR (Correct Alternation Ratio).

After extracting measurements from all tasks, we investigated whether they were correlated with each other, which would indicate that skills to perform correlated tasks could be similar. We first investigated if there was any correlation between the open field and novel object recognition, which are exploratory, with the y-maze, which specifically tests working memory. As expected, we found that a high number of arm entries correlated with a high distance moved in the Y-maze (r = 0.96, p = 2.0-e04). We also found that, although not correlated with object exploration time ratio, a higher number of arm entries in the Y-maze correlated with less time exploring the novel object in the novel object recognition (r = −0.686, p = 4.6e-02; Fig 2C-E). This correlation might indicate that mice exhibiting higher escape responses would have higher number of arm entries and be less likely to explore the novel object during the NOR task.

Correlation analysis between each measure from the behavior tasks and the vertical pole task indicated significant correlation between the traveled distance in novel object recognition and the two measures time to descend and time to turn in the vertical pole task (Fig 2D). A greater traveled distance in the novel object recognition task correlated with a quicker descent (r = −0.817, p = 1.0e-02; Fig 2F), and a longer time to turn (r = 0.717, p = 3.6e-02; Fig 2G) in the vertical pole task. This finding suggests that animals with a greater traveled distance in the novel object recognition task will perform well in the descent, but at the same time take longer to turn.

When evaluating the correlation between all measurements from all behavioral tasks and the pellet reaching task measures (Fig 3A), we found only one strong correlation. No correlations between vertical pole and pellet reaching variables was found (abs. r *<* −0.588; p *>* 0.074; Fig 3B-E), and a positive correlation is shown between the time spent in center in the open field test and the success ratio in the pellet reaching task (r = 0.661, p = 4.6e-02; Fig 3F). Interestingly, the analysis displayed a positive correlation between the pellet reaching success ratio and the Y-maze correct alternation ratio. Individuals who had the correct pattern when making a turn in the Y-maze (indicating better working memory) had a higher success ratio when reaching for sugar pellets (r = 0.93, p = 8.0e-04; Fig 3G).

**Figure 3:**
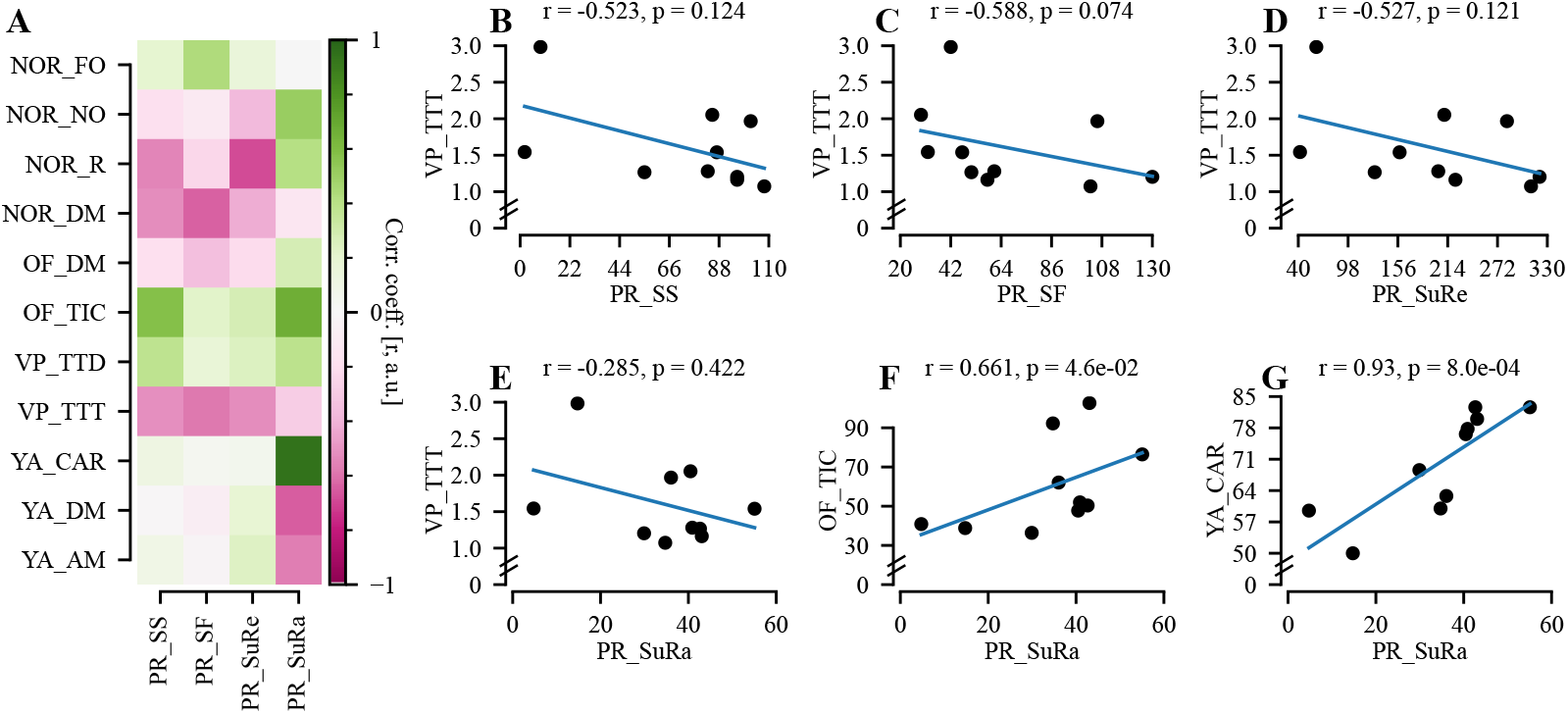
Y-Maze measures correlate with pellet reaching measures. A) Correlation matrix between each task measure versus the pellet reaching task measures. B-E) Correlation between vertical pole and pellet reaching measures. F) Open field time in center showed a positive correlation with the pellet reaching success ratio. G) Y-maze correct alternations showed a positive correlation with the pellet reaching success ratio. Abbreviations: NOR (Novel Object Recognition); OF (Open Field); VP (Vertical Pole); YA (Y-maze); DM (Distance Moved); FO (Familiar Object); NO (Novel Object); R (Ratio); TIC (Time In Center); TTD (Time To Descend); TTT (Time To Turn); AM (Alternations Maximum - total number of possible alternations); CAR (Correct Alternation Ratio); PR (Pellet Reaching); SS (Sum of Success); SF (Sum of Fail); SuRe (Sum of all Reaches); SuRa (Success Ratio).

To evaluate whether performance in the Y-maze predicts performance in the pellet reaching task, we conducted a comprehensive predictive validity analysis using partial correlations, multiple linear regression, leave-one-out cross-validation (LOOCV), and bootstrap confidence intervals. Y-maze correct alternation ratio demonstrated a very strong association with pellet reaching success ratio (partial Spearman *ρ* = 0.93, p = 8.0e-04; Fig 4A-B, blue), explaining 66.6% of the variance (*R*^2^ = 0.666, 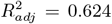; Fig 4C-D, blue). The model showed robust out-of-sample predictive capability with a positive LOOCV (*R*^2^ = 0.489, LOO-RMSE = 9.947; Fig 4E, blue), and the bootstrap-derived 95% confidence interval for the Y-maze correct alternation ratio slope was narrow and well-separated from zero (CI = [0.439, 1.69]; Fig 4F, blue), indicating stable predictive effects.

**Figure 4:**
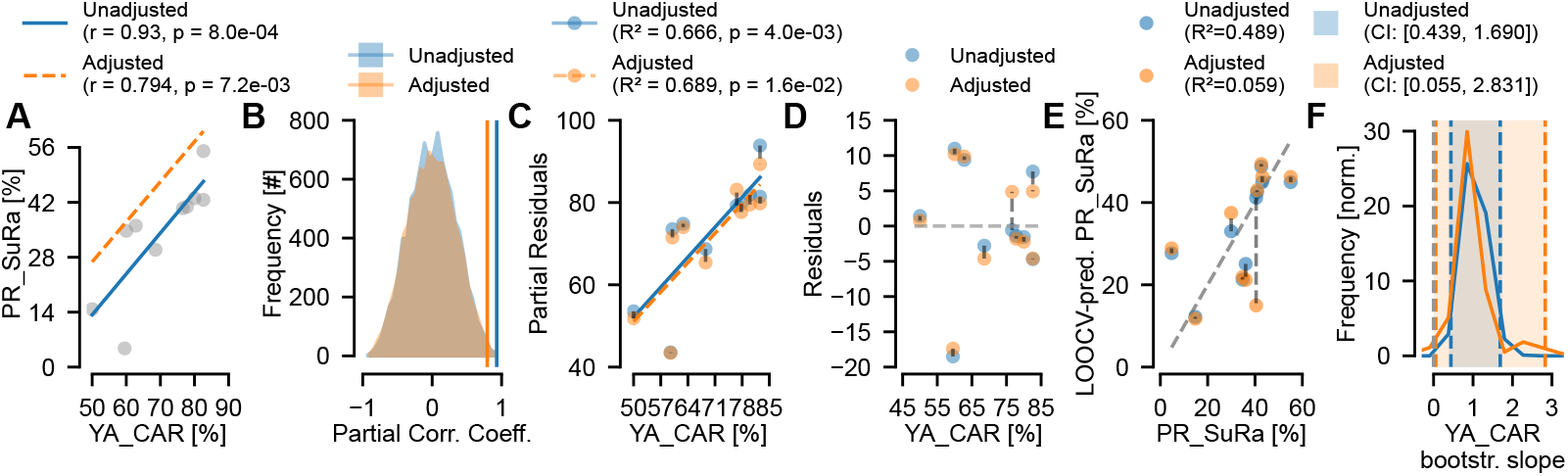
Y-maze performance robustly predicts pellet reaching performance in both unadjusted (blue) and adjusted (orange) models. A) Scatter plot of Y-Maze correct alternation ratio (YA CAR) versus pellet reaching success (PR SuRa) with regression lines. B) Distribution of permutation null distributions for partial Spearman correlation coefficients. The vertical lines indicate the observed partial correlation coefficient. C) Partial residuals plot demonstrating the relationship between Y-Maze correct alternation ratio and residualized outcome after removing linear effects of distance moved and maximum alternations. Lines show the marginal slope of the predictor on the partial residuals. D) Residuals from regression models plotted against Y-Maze correct alternation ratio, showing deviation of observed outcomes from predicted values after accounting for confounders. The dashed gray line indicates zero residuals (perfect fit). E) Leave-one-out cross-validated (LOOCV) predictions of pellet reaching success against actual values, with the dashed gray line indicating perfect prediction. F) Distribution of bootstrap estimates of regression slope. Each density curve shows the empirical distribution of slope coefficients derived from resampling, with 95% confidence intervals (CI) shown as shaded regions. The vertical lines mark the lower and upper bounds of the confidence intervals.

When controlling for the Y-maze distance moved and total number of possible alternations, the predictive strength remained statistically significant but was attenuated, consistent with partial mediation by these covariates. The adjusted model still showed a strong association between y-maze correct alternation ratio and pellet reaching success ratio (partial Spearman correlation *ρ* = 0.794, p = 7.2e-03; Fig 4A-B, orange), with y-maze performance uniquely explaining 68.9% of the variance in pellet reaching performance after accounting for confounding factors 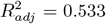 Fig 4C-D, orange). Despite the reduced effect size, the model maintained meaningful out-of-sample predictive validity with a positive LOOCV (*R*^2^ = 0.059, LOO-RMSE = 13.496; Fig 4E, orange). Critically, the bootstrap 95% confidence interval for the Y-maze correct alternation ratio slope in the corrected model remained entirely above zero (CI = [0.055, 2.831]; Fig 4F, orange), confirming that the predictive relationship persists even after rigorous adjustment for confounds. These findings collectively indicate that while the Y-maze distance moved and total number of possible alternations accounted for a portion of the observed association, Y-maze performance retains independent predictive utility for pellet reaching performance.

## Discussion

This study aimed to identify whether behavioral performance in memory and motor tasks correlates with subsequent motor learning success. By examining correlations between tasks such as the Y-maze and pellet reaching, we investigated potential pre-screening tasks that can be useful in future studies that require high motor learning capacity. The positive correlation between Y-maze performance and pellet reaching task success highlight the potential for contextual memory to enhance motor learning, as both processes rely on hippocampal function [35, 13, 36, 37]. This is further supported by the observation that motor learning requires coordinated activity between hippocampal and motor cortical networks, which work together to stabilize and adapt learned behaviors during memory consolidation [38]. These findings highlight the importance of contextual memory in motor learning tasks and suggest that the hippocampus interacts with motor functions in a task-dependent manner, influencing both the initial encoding of motor skills and their subsequent refinement.

Our findings show that the Y-maze task correlated with the ratio of successful reaches and that it can predict the performance in pellet reaching task. This suggests that the Y-maze might be an accurate choice for selecting candidates for a motor task study. However, it is important to note that the success ratio is not the only variable to consider when analyzing the results. For example, an individual who only tried to reach for a pellet twice and succeeded once would have a 50% success ratio. The inactivity or disinterest in the task makes it difficult to analyze the learning capacity of the individual if the only measure is the success ratio; hence, the total number of reaches is important.

The findings suggest that the relationship between the Y-maze correct alternation ratio and pellet reaching success ratio is not simply due to increased activity or trial-and-error behavior in the Y-maze. If this was the case, we would expect a correlation between these variables and the number of attempts in the pellet reaching task, reflecting a general tendency towards trial-and-error across both tests. However, our results indicate no such correlation exists, implying that the connection between the Y-maze correct alternation ratio and pellet reaching success is likely driven by other factors, such as specific cognitive processes or neural mechanisms, rather than mere activity levels. Further, we found that Y-maze correct alternation ratio is a reliable predictor of pellet reaching success ratio, even when accounting for distance moved and total number of alternations. This distinction is important because it highlights that the observed relationship is not a byproduct of general exploratory or motor activity, but may instead reflect more targeted aspects of learning or memory functions that are shared between the tasks.

Another intriguing finding reveals that animals that excelled in the pellet reaching task showed reduced stress levels by spending more time in the center of the open field arena, suggesting a correlation between decreasing anxiety and improved motor performance. This is similar to what is described in the literature where increased anxiety levels can have a negative impact on different perceptual-motor tasks [39]. Likewise, in the vertical pole, animals that spent less time descending from the pole tended to cover longer distances in the object recognition task, revealing an association between increased exploratory behavior, and motor coordination. Interestingly, no other behavior measure correlated with the pellet reaching task (Table 1, S1 Fig). One possible explanation to the findings that the vertical pole time to descend correlated with the total distance in the novel object recognition but not in the open field is that the open field task primarily measures locomotor activity and exploratory behavior in a more generalized context, while novel object recognition specifically evaluates recognition memory, which may involve more contextual and spatial processing that could also be relevant to the vertical pole task.

**Table 1:**
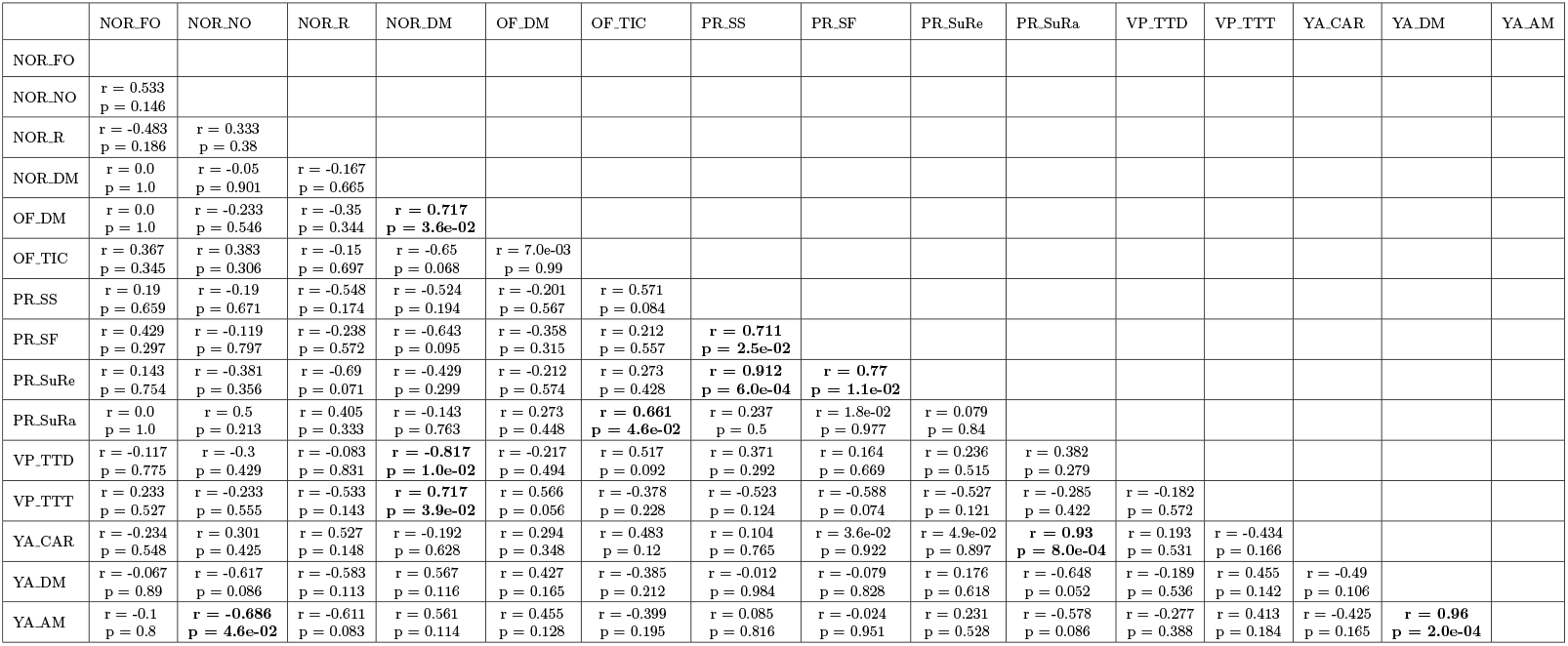
All possible correlations between task measures. The correlated measures are marked in bold. Abbreviations: NOR (Novel Object Recognition); OF (Open Field); PR (Pellet Reaching); VP (Vertical Pole); YA (Y-maze); FO (Familiar Object); NO (Novel Object); R (Ratio); DM (Distance Moved); TIC (Time In Center); SS (Sum of Success); SF (Sum of Fail); SuRe (Sum of all Reaches); SuRa (Success Ratio); TTD (Time To Descend); TTT (Time To Turn); CAR (Correct Alternation Ratio).

To reduce the risk of fatigue and stress affecting performance, all behavioral tests were limited to a maximum of 10 minutes per day. This approach minimized the likelihood that fatigue would significantly influence task outcomes. Additionally, our testing protocol relied on the animal’s natural innate rhythm, as it is not subjected to forced exhaustion methods such as forced swim test or fixed speeds in treadmills. This allows the animal to control its own performance in accordance with its individual pace and well-being, which may contribute to a reduction in stress levels. We also observed that although mice moved slower in the NOR test compared to the open field, they still covered higher distances at a reasonable speed, indicating that the reduced distance moved could be due to increased time in exploring the objects. Also, no significant differences in distance moved were found between open field (First test) and Y-Maze (third test), and we found no correlation between the distance moved in the open field and any pellet reaching measurement, indicating that the animals were not affected by fatigue on the subsequent test.

While our study provides valuable insights into the relationship between contextual memory and motor learning, it is important to acknowledge that several factors may influence the interpretation of results. One limitation is the relatively small sample size, which although powerful enough to find the correlations shown here, could not be powerful enough to detect more nuanced behavioral correlations. Another limitation is that, although literature confirms no sex-specific differences in the behavioral metrics used here for C57Cl6/J mice on the light phase of the light/dark cycle [19, 20, 21, 22], our use of female-only mice may limit the generalization of our results to both sexes. Although there is no difference in performance in C57BL/6 female mouse behavior due to the estrous cycle on exploratory measures [40], it could be important to consider the estrous phase in learning tasks since the influence of stress on reward learning is affected by the estrous cycle [41], and estrogen could modulate reward prediction errors and reinforce learning [42]. Future research should consider the potential impact of estrous cycle phases, thereby improving the generalizability and translational relevance of our results. Additionally, previous work confirms no significant differences between C57BL/6J male and female mice in subtle sensorimotor skills [ladder rung task; 19], spontaneous distance moved during the light-phase of the day cycle [20], novel object recognition exploration duration during the test trial [where a familiar object is replaced with a novel object; 21], or spatial memory performance on the Y-maze, except when under acute stress conditions [22]. Nevertheless, the potential for anxiety or locomotor activity and the changes in estrous cycle that can vary on the day of testing could introduce confounding variables, as these factors may influence performance in both memory and motor tasks by affecting motivation, exploration, or task engagement. To strengthen the findings, future studies should consider including both male and female mice, and implement a more comprehensive screening protocol to assess baseline anxiety prior to behavioral testing.

The use of mice and rats is very common in behavioral studies. When translating our findings from mice to rats, it is important to consider the distinct cognitive and motor abilities that exist between the two species. Although spatial memory in the y maze task is similarly affected in rats and mice upon Glutathione depletion [43], there are subtle differences in their learning and memory processes. For instance, a study using rats found that they spent more time exploring novel environments [44] without significant recognition memory differences compared to mice. Moreover, kinematic analysis showed that while both species can use a single forelimb to reach for food, rats exhibited better performance and more consistent results in tasks measuring reaching speed and accuracy [45]. Given these differences, it’s important to validate our findings using rat subjects before translating our results to this species, to ensure that the observed correlations between memory performance and motor learning are not specific to mice.

While multiple comparisons are commonly addressed through multiplicity corrections, these are primarily designed to control false positives under a global null hypothesis, such as “all pre-screening measures correlate with motor learning outcomes”. In exploratory studies where no broad inferences are made across multiple pairwise relationships, applying multiplicity correction to individual correlations do not appropriately adjust for Type I error rates, and instead inflate Type II errors by reducing statistical power, increasing the distance between observed correlations and their biological meaning. Given that our analysis focused on identifying specific, interpretable patterns rather than testing a comprehensive set of interrelationships among behavioral measures, we interpret the reported correlations as exploratory findings grounded in effect size (R) and contextual plausibility, rather than relying solely on p-values or multiplicity-adjusted thresholds.

The fixed order of tests performed in our study may introduce carryover effects, which could influence inter-task performance. However, our analysis focuses on correlations between spatial/memory tasks and a motor learning task, so our findings do not imply that performance in one task enhances another. Instead, the observed correlations - particularly between Y-maze and vertical pole with pellet reaching performance - likely reflect shared neural mechanisms, such as hippocampal and striatal involvement in both contextual memory and motor learning [35, 13, 36, 37]. While our conclusions are restricted to the independent relationship between performance in the tasks, not task-specific performance, we acknowledge that future studies should account for test order to control for these effects, as the observed patterns may be due to contextual memory interactions or task-dependent neural activation rather than direct practice effects, highlighting the potential for neural priming and cognitive transfer between memory and motor tasks.

## Conclusions

Results from this study are significant for future studies, especially behavioral studies using optogenetics or miniscopes, which involve invasive surgery that heavily rely on animals that can learn a task. Using a non-invasive pre-screening test enables researchers to only include animals likely to learn while minimizing the harm to non-learning mice. These results are particularly important for motor tasks like the pellet reaching task, which are time-consuming and involve food deprivation, potentially causing distress to the animals involved. A pre-selection process can therefore increase the effectiveness of subsequent behavioral studies, and at the same time, adhere to the principles of the 3 Rs - replacement, reduction, and refinement by reducing the number of animals used in the experiments, as non-learners are not included. In summary, we conclude that the Y-maze task is a suitable predictor test to pre-select good candidates for behavioral studies that include motor skill acquisition.

## Author Contributions

Conceptualization (KK, SS, BC), Methodology (IK, SS, BC), Validation (SS, BC), Resources (KK), Data curation (IK, SS, BC), Formal analysis (TM), Visualization (IK, TM, BC), Supervision (TM, KK, SS, BC), Original draft preparation (IK, BC), Writing - Reviewing and editing (IK, TM, KK, SS, BC).

## Acknowledgments

We thank Uppsala University behavioral facility (UUBF) for support.

## Data Availability Statement

The datasets generated and/or analyzed in the current study will be publicly made available (Zenodo, DOI to be provided upon acceptance).

## References

[1] Yadin Dudai, Avi Karni, and Jan Born. The consolidation and transformation of memory. Neuron, 88(1):20–32, October 2015. ISSN 0896-6273. doi: 10.1016/j.neuron.2015.09.004. URL http://dx.doi.org/10.1016/j.neuron.2015.09.004.

[2] Emily T. Cowan, Anna C. Schapiro, Joseph E. Dunsmoor, and Vishnu P. Murty. Memory consolidation as an adaptive process. Psychonomic Bulletin & Review, 28(6):1796–1810, July 2021. ISSN 1531-5320. doi: 10.3758/s13423-021-01978-x. URL http://dx.doi.org/10.3758/s13423-021-01978-x.

[3] Cathrine V. Jansson-Boyd and Peter Bright. Memory and learning, pages 93–118. Elsevier, 2024. ISBN 9780443135811. doi: 10.1016/b978-0-443-13581-1.00006-6. URL http://dx.doi.org/10.1016/b978-0-443-13581-1.00006-6.

[4] Daniel B. Willingham. A neuropsychological theory of motor skill learning. Psychological Review, 105(3):558–584, 1998. doi: 10.1037/0033-295x.105.3.558. URL https://doi.org/10.1037/0033-295x.105.3.558.

[5] Eran Dayan and Leonardo G. Cohen. Neuroplasticity subserving motor skill learning. Neuron, 72(3):443–454, November 2011. ISSN 0896-6273. doi: 10.1016/j.neuron.2011.10.008. URL http://dx.doi.org/10.1016/j.neuron.2011.10.008.

[6] Julien Doyon, Pierre Bellec, Rhonda Amsel, Virginia Penhune, Oury Monchi, Julie Carrier, Stéphane Lehéricy, and Habib Benali. Contributions of the basal ganglia and functionally related brain structures to motor learning. Behavioural Brain Research, 199(1):61–75, April 2009. ISSN 0166-4328. doi: 10.1016/j.bbr.2008.11.012. URL http://dx.doi.org/10.1016/j.bbr.2008.11.012.

[7] Risa Kawai, Timothy Markman, Rajesh Poddar, Raymond Ko, Antoniu L. Fantana, Ashesh K. Dhawale, Adam R. Kampff, and Bence P. Ã–lveczky. Motor cortex is required for learning but not for executing a motor skill. Neuron, 86(3):800– 812, may 2015. doi: 10.1016/j.neuron.2015.03.024. URL https://doi.org/10.1016/j.neuron.2015.03.024.

[8] Chris I. De Zeeuw and Michiel M. Ten Brinke. Motor learning and the cerebellum. Cold Spring Harbor Perspectives in Biology, 7(9):a021683, September 2015. ISSN 1943-0264. doi: 10.1101/cshperspect.a021683. URL http://dx.doi.org/10.1101/cshperspect.a021683.

[9] Darby M. Losey, Jay A. Hennig, Emily R. Oby, Matthew D. Golub, Patrick T. Sadtler, Kristin M. Quick, Stephen I. Ryu, Elizabeth C. Tyler-Kabara, Aaron P. Batista, Byron M. Yu, and Steven M. Chase. Learning leaves a memory trace in motor cortex. Current Biology, 34(7):1519–1531.e4, April 2024. ISSN 0960-9822. doi: 10.1016/j.cub.2024.03.003. URL http://dx.doi.org/10.1016/j.cub.2024.03.003.

[10] David I. Anderson, Keith R. Lohse, Thiago Costa Videira Lopes, and A. Mark Williams. Individual differences in motor skill learning: Past, present and future. Human Movement Science, 78:102818, August 2021. ISSN 0167-9457. doi: 10.1016/j.humov.2021.102818. URL http://dx.doi.org/10.1016/j.humov.2021.102818.

[11] Sally Kelliny, Liying Lin, Isaac Deng, Jing Xiong, Fiona Zhou, Mohammed Al-Hawwas, Larisa Bobrovskaya, and XinFu Zhou. A new approach to model sporadic alzheimer’s disease by intracerebroventricular streptozotocin injection in app/ps1 mice. Molecular Neurobiology, 58(8):3692–3711, April 2021. ISSN 1559-1182. doi: 10.1007/s12035-021-02338-5. URL http://dx.doi.org/10.1007/s12035-021-02338-5.

[12] Stan B. Floresco, Jeremy K. Seamans, and Anthony G. Phillips. Selective roles for hippocampal, prefrontal cortical, and ventral striatal circuits in radial-arm maze tasks with or without a delay. The Journal of Neuroscience, 17(5): 1880–1890, 3 1997. doi: 10.1523/jneurosci.17-05-01880.1997. URL https://doi.org/10.1523/jneurosci.17-05-01880.1997.

[13] Tiaotiao Liu, Wenwen Bai, Mi Xia, and Xin Tian. Directional hippocampal-prefrontal interactions during working memory. Behavioural Brain Research, 338:1–8, February 2018. ISSN 0166-4328. doi: 10.1016/j.bbr.2017.10.003. URL http://dx.doi.org/10.1016/j.bbr.2017.10.003.

[14] Joo-Hyun Song. The role of attention in motor control and learning. Current Opinion in Psychology, 29:261–265, October 2019. ISSN 2352-250X. doi: 10.1016/j.copsyc.2019.08.002. URL http://dx.doi.org/10.1016/j.copsyc.2019.08.002.

[15] Ian Q Whishaw, Paul Whishaw, and Bogdan Gorny. The structure of skilled forelimb reaching in the rat: A movement rating scale. Journal of Visualized Experiments, (18), August 2008. ISSN 1940-087X. doi: 10.3791/816. URL http://dx.doi.org/10.3791/816.

[16] Richardson N Leão, Sanja Mikulovic, Katarina E Leão, Her-many Munguba, Henrik Gezelius, Anders Enjin, Kalicharan Patra, Anders Eriksson, Leslie M Loew, Adriano B L Tort, and Klas Kullander. OLM interneurons differentially modulate CA3 and entorhinal inputs to hippocampal CA1 neurons. Nature Neuroscience, 15(11):1524–1530, oct 2012. doi: 10.1038/nn.3235. URL https://doi.org/10.1038/nn.3235.

[17] Samer Siwani, Arthur S.C. França, Sanja Mikulovic, Amilcar Reis, Markus M. Hilscher, Steven J. Edwards, Richardson N. Leão, Adriano B.L. Tort, and Klas Kullander. OLMα2 cells bidirectionally modulate learning. Neuron, 99(2):404–412.e3, July 2018. ISSN 0896-6273. doi: 10.1016/j.neuron.2018.06.022. URL http://dx.doi.org/10.1016/j.neuron.2018.06.022.

[18] Thawann Malfatti, Anna Velica, Jéssica Winne, Barbara Ciralli, Katharina Henriksson, George Nascimento, Richardson Leao, and Klas Kullander. Increased layer 5 martinotti cell excitation reduces pyramidal cell population plasticity and improves learned motor execution. November 2025. doi: 10.7554/elife.109286.1. URL http://dx.doi.org/10.7554/eLife.109286.1.

[19] Michael R. Hunsaker, Ramona E. von Leden, Binh T. Ta, Naomi J. Goodrich-Hunsaker, Gloria Arque, Kyoungmi Kim, Rob Willemsen, and Robert F. Berman. Motor deficits on a ladder rung task in male and female adolescent and adult cgg knock-in mice. Behavioural Brain Research, 222(1):117–121, September 2011. ISSN 0166-4328. doi: 10.1016/j.bbr.2011.03.039. URL http://dx.doi.org/10.1016/j.bbr.2011.03.039.

[20] Briana J. Bernstein, Dalisa R. Kendricks, Sydney Fry, Leslie Wilson, Bastijn Koopmans, Maarten Loos, Korey D. Stevanovic, and Jesse D. Cushman. Sex differences in spontaneous behavior and cognition in mice using an automated behavior monitoring system. Physiology & Behavior, 283:114595, September 2024. ISSN 0031-9384. doi: 10.1016/j.physbeh.2024.114595. URL http://dx.doi.org/10.1016/j.physbeh.2024.114595.

[21] Charles J. Heyser, Caitlyn H. McNaughton, Donna Vishnevetsky, and Allen A. Fienberg. Methylphenidate restores novel object recognition in darpp-32 knockout mice. Behavioural Brain Research, 253:266–273, September 2013. ISSN 0166-4328. doi: 10.1016/j.bbr.2013.07.031. URL http://dx.doi.org/10.1016/j.bbr.2013.07.031.

[22] Cheryl D Conrad, Jamie L Jackson, Lindsay Wieczorek, Sarah E Baran, James S Harman, Ryan L Wright, and Donna L Korol. Acute stress impairs spatial memory in male but not female rats: influence of estrous cycle. Pharmacology Biochemistry and Behavior, 78(3):569–579, July 2004. ISSN 0091-3057. doi: 10.1016/j.pbb.2004.04.025. URL http://dx.doi.org/10.1016/j.pbb.2004.04.025.

[23] Robert Lalonde. The neurobiological basis of spontaneous alternation. Neuroscience & Biobehavioral Reviews, 26(1): 91–104, 2002.

[24] Ian N Krout, Alexandria White, and Tim Sampson. Pole test assessment v1. January 2024. doi: 10.17504/protocols.io.5qpvo34k9v4o/v1. URL http://dx.doi.org/10.17504/protocols.io.5qpvo34k9v4o/v1.

[25] Chia-Chien Chen, Anthony Gilmore, and Yi Zuo. Study motor skill learning by single-pellet reaching tasks in mice. Journal of Visualized Experiments, (85), March 2014. ISSN 1940-087X. doi: 10.3791/51238. URL http://dx.doi.org/10.3791/51238.

[26] Satoshi Manita, Koji Ikezoe, and Kazuo Kitamura. A novel device of reaching, grasping, and retrieving task for head-fixed mice. Frontiers in Neural Circuits, 16, May 2022. ISSN 1662-5110. doi: 10.3389/fncir.2022.842748. URL http://dx.doi.org/10.3389/fncir.2022.842748.

[27] Miguel A. Garcća-Pérez. Use and misuse of corrections for multiple testing. Methods in Psychology, 8:100120, November 2023. ISSN 2590-2601. doi: 10.1016/j.metip.2023.100120. URL http://dx.doi.org/10.1016/j.metip.2023.100120.

[28] Pauli Virtanen, Ralf Gommers, Travis E. Oliphant, Matt Haberland, Tyler Reddy, David Cournapeau, Evgeni Burovski, Pearu Peterson, Warren Weckesser, Jonathan Bright, Stéfan J. van der Walt, Matthew Brett, Joshua Wilson, K. Jarrod Millman, Nikolay Mayorov, Andrew R. J. Nelson, Eric Jones, Robert Kern, Eric Larson, C J Carey, Ilhan Polat, Yu Feng, Eric W. Moore, Jake VanderPlas, Denis Laxalde, Josef Perktold, Robert Cimrman, Ian Henriksen, E. A. Quintero, Charles R. Harris, Anne M. Archibald, Antônio H. Ribeiro, Fabian Pedregosa, Paul van Mulbregt, and SciPy 1.0 Contributors. SciPy 1.0: Fundamental Algorithms for Scientific Computing in Python. Nature Methods, 17:261–272, 2020. doi: 10.1038/s41592-019-0686-2.

[29] Charles R. Harris, K. Jarrod Millman, Stéfan J van der Walt, Ralf Gommers, Pauli Virtanen, David Cournapeau, Eric Wieser, Julian Taylor, Sebastian Berg, Nathaniel J. Smith, Robert Kern, Matti Picus, Stephan Hoyer, Marten H. van Kerkwijk, Matthew Brett, Allan Haldane, Jaime Fernández del Rćo, Mark Wiebe, Pearu Peterson, Pierre Gérard-Marchant, Kevin Sheppard, Tyler Reddy, Warren Weckesser, Hameer Abbasi, Christoph Gohlke, and Travis E. Oliphant. Array programming with NumPy. Nature, 585:357–362, 2020. doi: 10.1038/s41586-020-2649-2.

[30] Thawann Malfatti and Barbara Ciralli. Sciscripts: a python library for controlling devices, running experiments and analyzing data, November 2023. URL https://doi.org/10.5281/zenodo.4045872.

[31] John D. Hunter. Matplotlib: A 2d Graphics Environment. Computing in Science & Engineering, 9(3):90–95, 2007. ISSN 1521-9615. doi: 10.1109/MCSE.2007.55. URL http://ieeexplore.ieee.org/document/4160265/.

[32] Inkscape Project. Inkscape, 2022. URL https://inkscape.org. Software.

[33] Thawann Malfatti and Barbara Ciralli. Prtbehpred: Performance correlation between pellet-reaching task and memory/motor tasks, October 2025. URL https://doi.org/10.5281/zenodo.17484469.

[34] Marianne Leger, Anne Quiedeville, Valentine Bouet, Benoît Haelewyn, Michel Boulouard, Pascale Schumann-Bard, and Thomas Freret. Object recognition test in mice. Nature Protocols, 8(12):2531–2537, November 2013. ISSN 1750-2799. doi: 10.1038/nprot.2013.155. URL http://dx.doi.org/10.1038/nprot.2013.155.

[35] Geneviève Albouy, Bradley R. King, Pierre Maquet, and Julien Doyon. Hippocampus and striatum: Dynamics and interaction during acquisition and sleep-related motor sequence memory consolidation. Hippocampus, 23(11):985–1004, October 2013. ISSN 1098-1063. doi: 10.1002/hipo.22183. URL http://dx.doi.org/10.1002/hipo.22183.

[36] Fabian Chersi and Neil Burgess. The cognitive architecture of spatial navigation: Hippocampal and striatal contributions. Neuron, 88(1):64–77, October 2015. ISSN 0896-6273. doi: 10.1016/j.neuron.2015.09.021. URL http://dx.doi.org/10.1016/j.neuron.2015.09.021.

[37] Arnaud Boutin, Basile Pinsard, Arnaud Boré, Julie Carrier, Stuart M. Fogel, and Julien Doyon. Transient synchronization of hippocampo-striato-thalamo-cortical networks during sleep spindle oscillations induces motor memory consolidation. NeuroImage, 169:419–430, April 2018. ISSN 1053-8119. doi: 10.1016/j.neuroimage.2017.12.066. URL http://dx.doi.org/10.1016/j.neuroimage.2017.12.066.

[38] Jaekyung Kim, Abhilasha Joshi, Loren Frank, and Karunesh Ganguly. Cortical–hippocampal coupling during manifold exploration in motor cortex. Nature, 613(7942):103–110, December 2022. ISSN 1476-4687. doi: 10.1038/s41586-022-05533-z. URL http://dx.doi.org/10.1038/s41586-022-05533-z.

[39] Arne Nieuwenhuys and Raôul RD Oudejans. Anxiety and performance: perceptual-motor behavior in high-pressure contexts. Current Opinion in Psychology, 16:28–33, August 2017. ISSN 2352-250X. doi: 10.1016/j.copsyc.2017.03.019. URL http://dx.doi.org/10.1016/j.copsyc.2017.03.019.

[40] Pei-Yun Zeng, Ya-Hsuan Tsai, Chih-Lin Lee, Yu-Kai Ma, and Tsung-Han Kuo. Minimal influence of estrous cycle on studies of female mouse behaviors. Frontiers in Molecular Neuroscience, 16, July 2023. ISSN 1662-5099. doi: 10.3389/fnmol.2023.1146109. URL http://dx.doi.org/10.3389/fnmol.2023.1146109.

[41] Morgan P. Johnston, Brandon I. Garcia-Castañeda, Leonor G. Cedillo, Sachi K. Patel, Victoria S. Vargas, and Matthew J. Wanat. Estrous cycle stage gates the effect of stress on reward learning. Neuropsychopharmacology, 51(3):650–660, July 2025. ISSN 1740-634X. doi: 10.1038/s41386-025-02170-8. URL http://dx.doi.org/10.1038/s41386-025-02170-8.

[42] Carla E. M. Golden, Audrey C. Martin, Daljit Kaur, Andrew Mah, Diana H. Levy, Takashi Yamaguchi, Amy W. Lasek, Dayu Lin, Chiye Aoki, and Christine M. Constantinople. Estrogen modulates reward prediction errors and reinforcement learning. Nature Neuroscience, 28(12):2502â€”2514, November 2025. ISSN 1546-1726. doi: 10.1038/s41593-025-02104-z. URL http://dx.doi.org/10.1038/s41593-025-02104-z.

[43] Olivia Dean, Ashley I. Bush, Michael Berk, David L. Copolov, and Maarten van den Buuse. Glutathione depletion in the brain disrupts short-term spatial memory in the y-maze in rats and mice. Behavioural Brain Research, 198(1):258–262, March 2009. ISSN 0166-4328. doi: 10.1016/j.bbr.2008.11.017. URL http://dx.doi.org/10.1016/j.bbr.2008.11.017.

[44] Alexis M. Stranahan. Similarities and differences in spatial learning and object recognition between young male c57bl/6j mice and sprague-dawley rats. Behavioral Neuroscience, 125 (5):791–795, 2011. ISSN 0735-7044. doi: 10.1037/a0025133. URL http://dx.doi.org/10.1037/a0025133.

[45] Ian Q. Whishaw. An endpoint, descriptive, and kinematic comparison of skilled reaching in mice (mus musculus) with rats (rattus norvegicus). Behavioural Brain Research, 78(2): 101–111, 8 1996. doi: 10.1016/0166-4328(95)00236-7. URL https://doi.org/10.1016/0166-4328(95)00236-7.

