## Supplemental File for "Y-maze performance predicts refined motor learning in mice"

### Supplementary figure

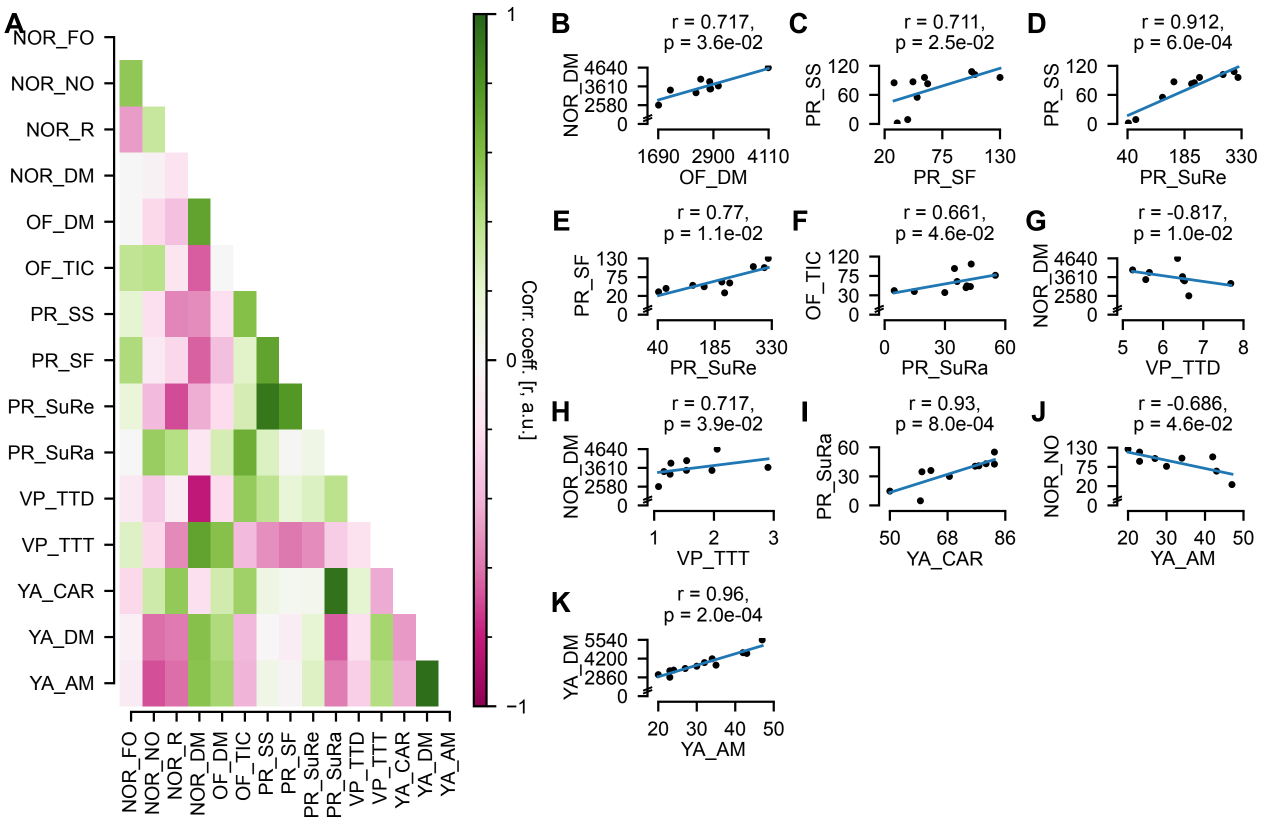


**Supplementary Figure 1:** Matrix of all possible correlations in the tasks (A) and plots of all measures that are correlated (B-K). Abbreviations: NOR (Novel Object Recognition); OF (Open Field); PR (Pellet Reaching); VP (Vertical Pole); YA (Y-maze); FO (Familiar Object); NO (Novel Object); R (Ratio); DM (Distance Moved); TIC (Time In Center); SS (Sum of Success); SF (Sum of Fail); SuRe (Sum of all Reaches); SuRa (Success Ratio); TTD (Time To Descend); TTT (Time To Turn); AM (Alternations Maximum - total number of possible alternations); CAR (Correct Alternation Ratio).
